# Oscillatory Hypoxia Induced Unfolded Protein Folding Response Gene Expression Predicts Low Survival in Human Breast Cancer Patients

**DOI:** 10.1101/2024.01.25.577274

**Authors:** Yasir Suhail, Yamin Liu, Wenqiang Du, Junaid Afzal, Xihua Qiu, Amina Atiq, Paola Vera-Licona, Eran Agmon, Kshitiz

## Abstract

Hypoxia is one of the key factors in the tumor microenvironment regulating nearly all steps in the metastatic cascade in many cancers, including in breast cancer. The hypoxic regions can however be dynamic with the availability of oxygen fluctuating or oscillating. The canonical response to hypoxia is relayed by transcription factor HIF-1, which is stabilized in hypoxia and acts as the master regulator of a large number of downstream genes. However, HIF-1 transcriptional activity can also fluctuate either due to unstable hypoxia, or by lactate mediated non-canonical degradation of HIF-1. Our understanding of how oscillatory hypoxia or HIF-1 activity specifically influence cancer malignancy is very limited. Here, using MDA-MB-231 cells as a model of triple negative breast cancer characterized by severe hypoxia, we measured the gene expression changes induced specifically by oscillatory hypoxia. We found that oscillatory hypoxia can specifically regulate gene expression differently, and at times opposite to stable hypoxia. Using The Cancer Genome Atlas (TCGA) RNAseq data of human cancer samples, we show that the oscillatory specific gene expression signature in MDA-MB-231 is enriched in most human cancers, and prognosticate low survival in breast cancer patients. In particular, we found that oscillatory hypoxia, unlike stable hypoxia, induces unfolded protein folding response (UPR) in cells resulting in gene expression predicting reduced survival.

## Introduction

As tumors expand and grow, they can outstrip their nutrient supply locally to create regions of hypoxia. Hypoxia acts as a key microenvironmental factor influencing nearly all steps in the metastatic cascade. The chief regulator of hypoxic response in cells is the transcription factor Hypoxia-Inducible Factor 1 (HIF-1), which is acutely sensitive to lack of oxygen, and can rapidly induce transcription of a large number of downstream genes with the onset of hypoxia^1^. HIF-1 is a heterodimeric transcription factor (TF) composed of two subunits, with HIF-1α being the regulated unit. HIF-1α is constantly expressed, transcribed, and is rapidly degraded in an oxygen dependent manner by ubiquitin mediated proteasomal degradation^2^. Lack of oxygen can stabilize HIF-1α, which then binds to HIF-1β. The composite molecule, HIF-1 is a TF which binds to the HIF responsive element (HRE), and acts as a master regulator of a large gene-set affecting angiogenesis, metabolic transition, cell migration, immunogenicity, drug resistance, and other cancer-related phenotypes (**Figure 1A**)^3^.

**Figure 1.**
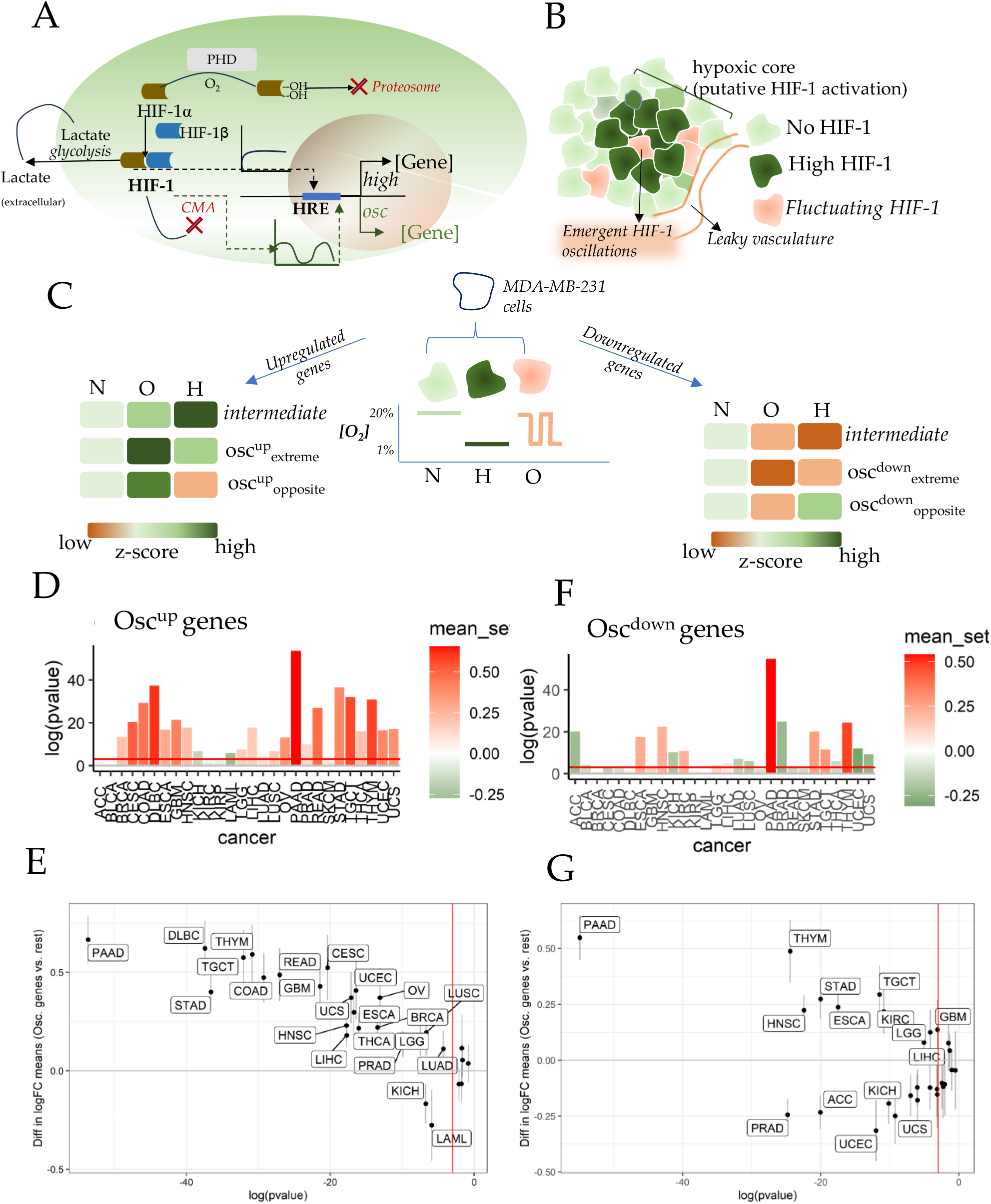
Hypoxic oscillation specific genes are enriched in human cancers. (A) Schematic showing the dynamics of HIF1a in hypoxic oscillations. (B) The population of HIF1a oscillatory cells in the tumor core. (C) Certain oscillation specific genes show either a larger differential expression in hypoxic oscillations compared to stable hypoxia, while others show differential expression in the opposite direction to stable hypoxia. (D-E) The average expression of oscillatory expression genes in different cancer samples compared to the matched normal tissue for genes up-regulated in oscillatory hypoxia (D) and those down-regulated in hypoxic oscillations (E).

However, recent years have revealed that many regions in cancers do not exhibit stable, and can exhibit “intermittent/fluctuating” hypoxia^4,5^. Oxygen availability in these localized regions is not stable arising from many potential mechanisms, including poor and leaky vasculature, vascular mimicry by endothelial transformation of epithelial or mesenchymal cells, as well as other mechanisms like changes in temperature^5,6^. Intermittent hypoxia can be more systemic, for example, from obstructive sleep apnea affecting millions of people (**Figure 1B**). However, HIF-1 stabilization in hypoxia itself could be dysregulated, resulting in endogenous oscillations in the transcriptional activity of HIF-1^7,8^. We had found that lactate, a byproduct of HIF-1 induced glycolysis in cancer cells, can prime a subpopulation to undergo oscillations in HIF-1 transcriptional activity. These oscillations were achieved by lactate mediated degradation of HIF-1 by chaperone mediated autophagy (CMA), a non-canonical mechanism regulating HIF-1 abundance (**Figure 1B**). Lactate can accumulate in very large amounts in cancers, reaching as high as 40mM, and therefore it can act as a key microenvironmental factor in the tumor microenvironment^9^. In addition, other factors, including reactive oxygen species, can both stabilize and destabilize HIF-1 through different mechanisms^10^, providing parameter realms where HIF-1 levels can fluctuate (**Figure 1B**). Although instability, or fluctuations in hypoxia and HIF-1 levels have been described in recent years, its effect on cancer phenotypes is less well understood.

We sought to understand how oscillation in the hypoxic atmosphere can affect gene expression patterns in cancers differently from stable hypoxia. Using MDA-MB-231 cells as a model of triple negative breast cancer which can develop significant tumor hypoxia, we transcriptomically profiled cells in normoxia, hypoxia, and oscillatory hypoxia. We found that oscillation in hypoxia had a profound effect on gene expression. However, interestingly, as we had noted previously for HeLa cells^7^, we found genes which exhibited an “oscillatory specific signature” with gene expression opposite to stable hypoxia, or higher (or lower) than hypoxia. Examining the expression patterns of these genes across human cancers in The Cancer Genome Atlas (TCGA)^11^, we found that oscillatory hypoxia can significantly increase expression of genes enriched in many human cancers, and even reduce expression of genes which are negative enriched in human cancers. Furthermore, we also found that genes increased in oscillatory hypoxia prognosticate low survival. Our analysis highlights that heterogeneity in hypoxia, and HIF-1 activity can transform breast cancer cells to assume a more aggressive phenotype, both highlighting the importance of localized microenvironmental factors, as well as presenting opportunities to target development of phenotypic heterogeneity in breast cancers.

## Results

### Oscillatory hypoxia results in specific gene expression patterns enriched in many human cancers

Both emergent oscillations in HIF-1 abundance and activity, as well as oscillations in hypoxia are highly localized phenomena, affecting a relatively small number of cells within tumor. However, our report showed that these cells may be endowed with the property of escaping HIF-1 induced cell cycle arrest, and therefore can have an outsized contribution to cancer growth. Breast cancers are frequently hypoxic and can HIF-1 is activation is known to influence key aspects of breast cancer progression, including acquirement of drug resistance, metastasis, and modulation of extracellular matrix. To test the effect of how fluctuating or unstable hypoxia can affect breast cancer cells differently from stable hypoxia, we cultured MDA-MB-231 cells on oxygen permeable surfaces. Cells were subjected to ambient oxygen, 1% O_2_, and an oscillatory regimen of hypoxia (1% O_2_ for 60 min, followed by 20% O_2_ for 30 min) for 48 hours. RNA was immediately collected and sequenced for mRNA levels. Out of 17056 for which significant reads were sequenced, 6176 were differentially regulated between hypoxia and control, 3374 between oscillatory hypoxia and control, and 5137 between oscillatory and stable hypoxia. Our definition for differential regulation is a false discovery rate of 5% or less. We found that although most genes in oscillatory hypoxia (hyp^osc^) showed expression intermediately between normoxia, and hypoxia (hyp^hi^), as we had previously reported for HeLa cells^7^, there were a large number of genes with an unexpected pattern of expression in hyp^osc^. Many genes increased in expression in hyp^osc^ even more than in hyp^hi^ (Osc^up^extreme, 215 genes), while others increased in expression in hyp^osc^, although they had decreased in expression in stable hypoxia (Osc^up^opposite, 293 genes). Together, we refer to both gene-sets as oscillatory responsive genes (Osc genes). A similar set could be defined for genes that deceased in expression in hypoxia vs normoxia (Osc^down^, 470 genes) (Figure 1C). We then sought to test the relevance of genes that show an unexpected pattern of expression in oscillatory hypoxia in human cancers.

Towards this aim, we used The Human Cancer Genome Atlas (TCGA) database^11^ to test the enrichment of osc genes in cancer samples vs corresponding control tissue samples. We found that Osc^up^ genes in MDA-MB-231 cells were significantly enriched in most human cancers (**Figure 1D**). These included not just breast carcinomas (BRCA), but many other cancer types, including diffuse large B cell lymphoma (DLBCL), thymoma, and prostate cancer. These enrichment patterns sustained when we removed the systemic bias in transcriptomic analysis by calculating cancer vs normal expression differentials compared with the expression changes for all genes (**Figure 1E**). For nearly all cancers in the TCGA database, Osc^up^ genes were found to be enriched in cancers vs normal tissues.

The pattern in Osc^dn^ genes also showed a similarly anti-tumor pattern, although not as pronounced as in Osc^up^ genes. Osc^dn^ genes were significantly downregulated in many cancers, including ACC, PRAD, UCEC, THCA, and UCS, while also being upregulated in ESCA, HNSC, PAAD, and THYM. Removing systemic bias in transcriptomic analysis, we found that UCS, PRAD, ACC, KICH, and UCEC cancers showed a similar pattern of gene downregulation as observed in oscillatory hypoxia.

These data showed that overall, oscillatory hypoxia has a specific and a profound pro-tumor effect on cancer gene expression. Although hypoxia is considered to have a broadly pro-tumorigenic effect, HIF-1 activation, our data showed that fluctuations in hypoxia may induce gene expression changes which are even more associated with tumors across a broad range of human cancers.

### Oscillatory hypoxia induced gene expression is enriched in human breast cancer patients

We followed up with the original intent of investigating the effect of fluctuating hypoxia on breast cancer cells. Osc^up^ genes were significantly enriched in BRCA cancer tissues vs normal. Gene Set Enrichment Analysis (GSEA) showed that Osc^up^ genes significantly were enriched in BRCA genes when ranked in order of their differential expression between cancer and normal tissues (Figure 2A). We calculated the density of Osc^up^ genes vs all other genes in BRCA samples and found that osc^up^ genes are more up-regulated in cancer than by chance (**Figure 2B**).

**Figure 2.**
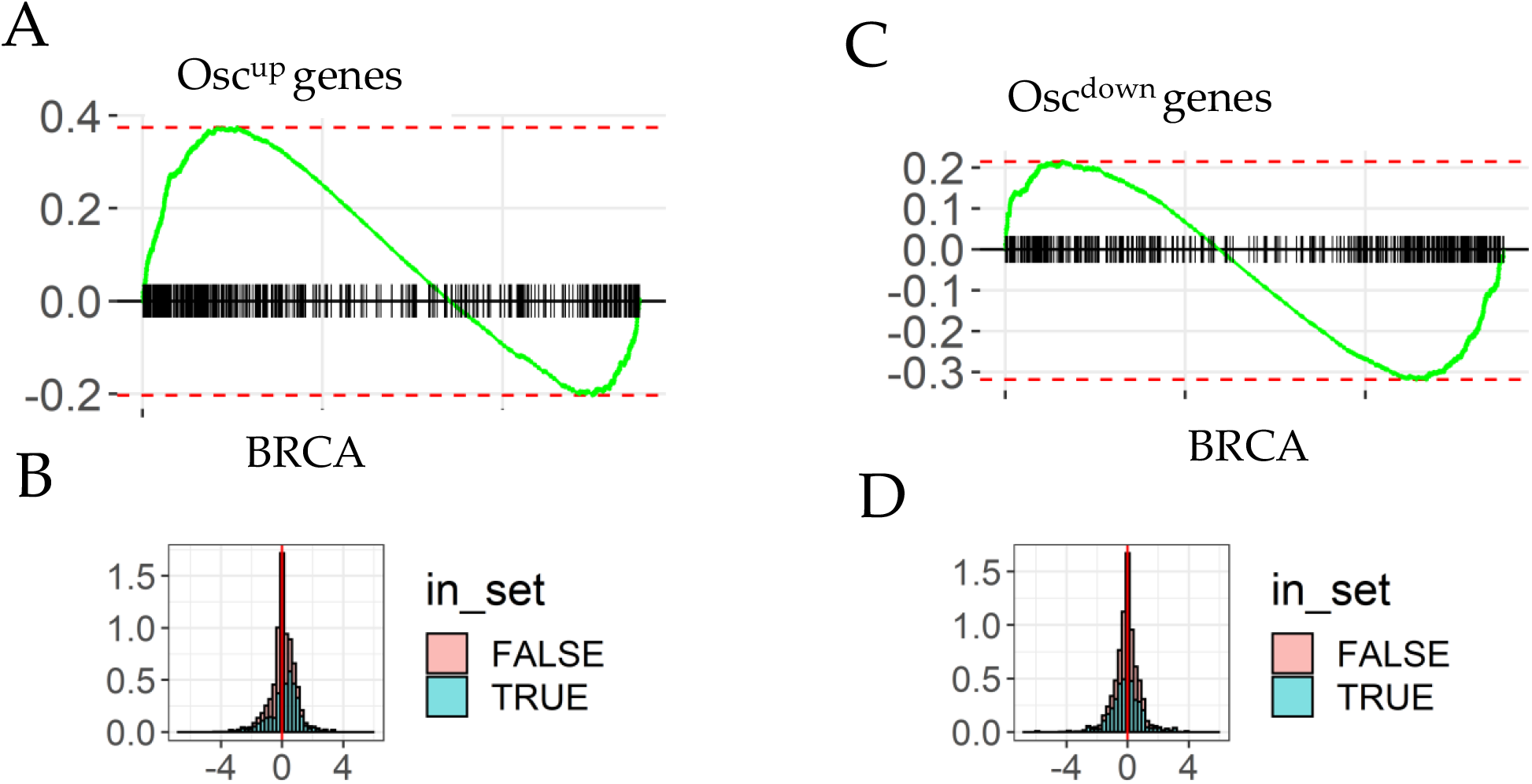
Hypoxic oscillation specific genes are differentially expressed in BRCA. GSEA plots showing enrichment among BRCA up-regulated genes for genes up-regulated in hypoxic oscillations (Osc^up^ genes, **A**) and for genes down-regulated in hypoxic oscillations (Osc^down^ genes, **C**). In each plot, the genes are ranked by differential expression in BRCA vs normal breast, and ticks on the x-axis correspond to Osc^up^ and Osc^down^ genes respectively. Histograms of the differential expression of all genes in BRCA vs normal breast, where the colors of the bars correspond to the genes belonging to Osc^up^ set (**B**), Osc^down^ set (**D**) or the rest of the genes.

**Figure 3.**
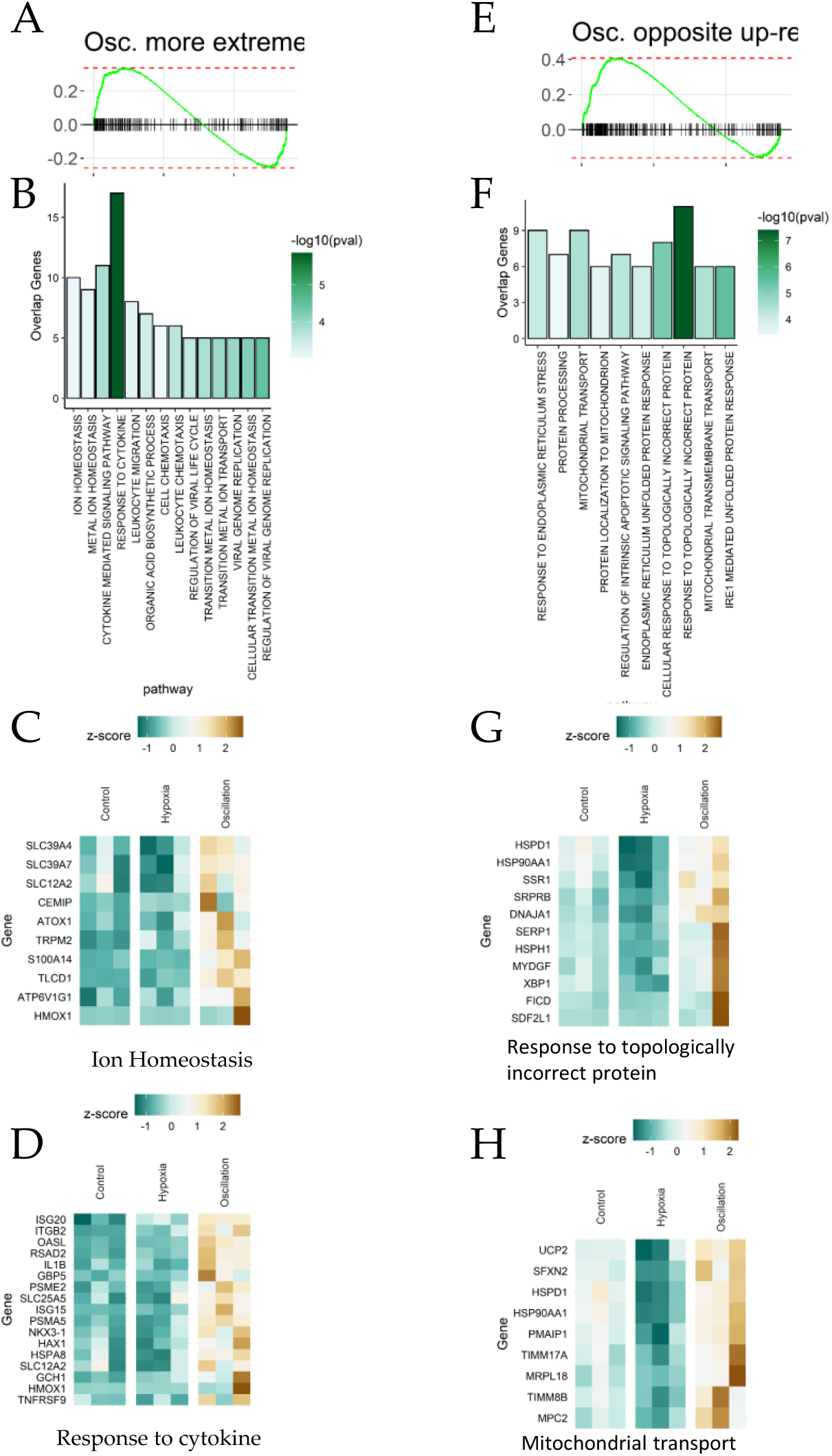
Hypoxic oscillation specific genes are selectively up-regulated in breast cancer and impact key pathways. (**A**) GSEA plot with oscillatory genes with higher gene expression than hypoxia (Osc_extreme_^up^), ranked in the order of BRCA vs normal gene expression. (**B**) Top GO pathways over-represented for genes in the leading edge of (**A**), i.e., genes that are both specifically up-regulated in oscillatory hypoxia and BRCA. (**C**) Heatmap of the genes overlapping with the leading edge of (**A**) and in the Ion Homeostasis pathway. (**D**) Overlapping genes in the Response to Cytokine GO pathway. (**D**) GSEA plot with oscillatory genes with gene expression opposite to hypoxia (Osc_opposite_^up^), ranked in the order of BRCA vs normal gene expression. (**E**) Top GO pathways over-represented for genes in the leading edge of (**E**), i.e., genes that are both specifically up-regulated in oscillatory hypoxia and BRCA. (**F**) Heatmap of the genes overlapping with the leading edge of (**E**) and in the Response to Topologically Incorrect Protein. (**F**) Overlapping genes in the Mitochondrial Transport GO pathway.

GSEA analysis of Osc^dn^ genes also showed an overall negative enrichment in BRCA cancers with a lower significance. Most Osc^dn^ genes were enriched in normal breast tissues vs cancer tissues, suggesting that emergent HIF-1 activity oscillations, or regions where hypoxia undergoes fluctuations can prime cancer cells to downregulate putative tumor-suppressor genes (**Figure 2C**). Histogram of the density of Osc^dn^ genes vs all other genes in BRCA samples confirmed the association (**Figure 2D**).

### Genes upregulated by hypoxic oscillations enriched in breast cancer patient samples indicate stress response

Genes upregulated in oscillatory hypoxia are enriched in many human cancers, including in breast cancer samples. We therefore sought to separately explore the gene set enrichment for Oscup genes, both those which were increased even more than in stable hypoxia (Osc^extreme^), as well those which showed a trend opposite to stable hypoxia (Osc^opposite^). Ranking genes according to their fold change between BRCA samples vs normal breast tissue controls, we performed GSEA analysis separately for either Osc^up^ gene-sets, finding enrichment of very different ontologies.

The top gene ontologies (GOs) in Osc^extreme^ enriched in BRCA were mostly related to ion homeostasis, and chemotaxis of leukocytes. Ion homeostasis is one of the lesser studied aspects of cancer metastasis, and has come into more careful investigation recently. Cancer metastasis is associated with changes in the homeostasis of various metal ions, including copper, sodium, calcium etc. as well as other ions including chloride. Ion channels, including aquaporins are known to regulate ameboidal migration in cancer cells by regulating water uptake^12^. A closer look at the top overlapping genes revealed several ion channels which were selectively upregulated in oscillatory hypoxia. These included SLC39 components responsible for Zn/Fe transport^13^, SLC12A2 encoding Na+ and Cl-absorption, as well as Ca2+ channels encoded by TRPM2. Leukocyte chemotaxis has been previously shown to be up-regulated in triple negative breast cancers. SenGupta et al^14^ found that tumor conditioned media from both triple negative breast cancer cell monolayers and spheroids recruit neutrophils. They also found that TNBC cells secrete neutrophil recruiting ligands more than ER+ breast cancer cells. Additionally, Ruffell et al^15^ found higher infiltration of leukocytes in breast cancer tumor tissue (including TNBC) compared to normal breast tissue.

In addition, we also found strong activation of GO associated with response to cytokines, revealing many genes with important role in cancer malignancy. These included many inflammation responsive genes including IL1B encoding Interleukin 1-beta, TNFRSF9 encoding a receptor of the TNF receptor superfamily, ISG15, and ISG20 encoding interferon stimulated exonucleases, as well as RSAD2 encoding another interferon stimulated antiviral protein.

Osc^up^ genes which showed expression opposite to that of hypoxia showed an even stronger enrichment in gene-set enriched in BRCA. This is remarkable, because expression of these genes suggests more complex transcriptional circuitry involving HIF-1 wherein these genes are expressed only when HIF-1 activity is dynamic/oscillatory, and not when it is stably high. We asked how fluctuations in O_2_ dosage change gene expression relevant to breast cancer differently from hypoxia. Similar to above, we identified GOs activated by the leading-edge genes obtained from the GSEA analysis with BRCA genes. We again surprisingly found that hypoxic oscillations resulted in activation of unfolded protein response (UPR) in MDA-MB-231 cells highlighted by multiple relevant GOs, including those associated with response to endoplasmic reticulum stress, protein processing, topologically incorrect protein, as well as IRE-1 mediated UPR. A closer exploration of the top genes in these GOs revealed those encoding chaperone proteins like HSPD1, HSPH1, as well as HSP90AA1. Another key GO upregulated was the related GO, mitochondrial transport, revealing above genes as well as UCP2 encoding uncoupling protein 2, a key mediator of oxidative response and mitochondrial membrane potential^16^.

We wanted to ask whether the Osc^up^ genes act together among patients. We computed the correlation between each pair of Osc^up^ genes, noticing a complex clustering pattern (**Figure 4A**). In order to gain insight into some specific genes, we cut the hierarchical clustering dendogram mid-way, we concentrate on 4 of the smaller clusters (**Figure 4B-E**). While the subcluster in **Figure 4B** mainly deals with unfolded protein response, the cluster in **Figure 4E** includes both unfolded protein response and translation related genes. The subclusters in **Figures 4C-D** are both immune related genes, with cluster in **Figure 4C** involved in T cell regulation while the one in **Figure 4D** related to viral response. It is possible that these groups of correlated genes are co-regulated by master regulators that are directly involved with HIF1A.

**Figure 4.**
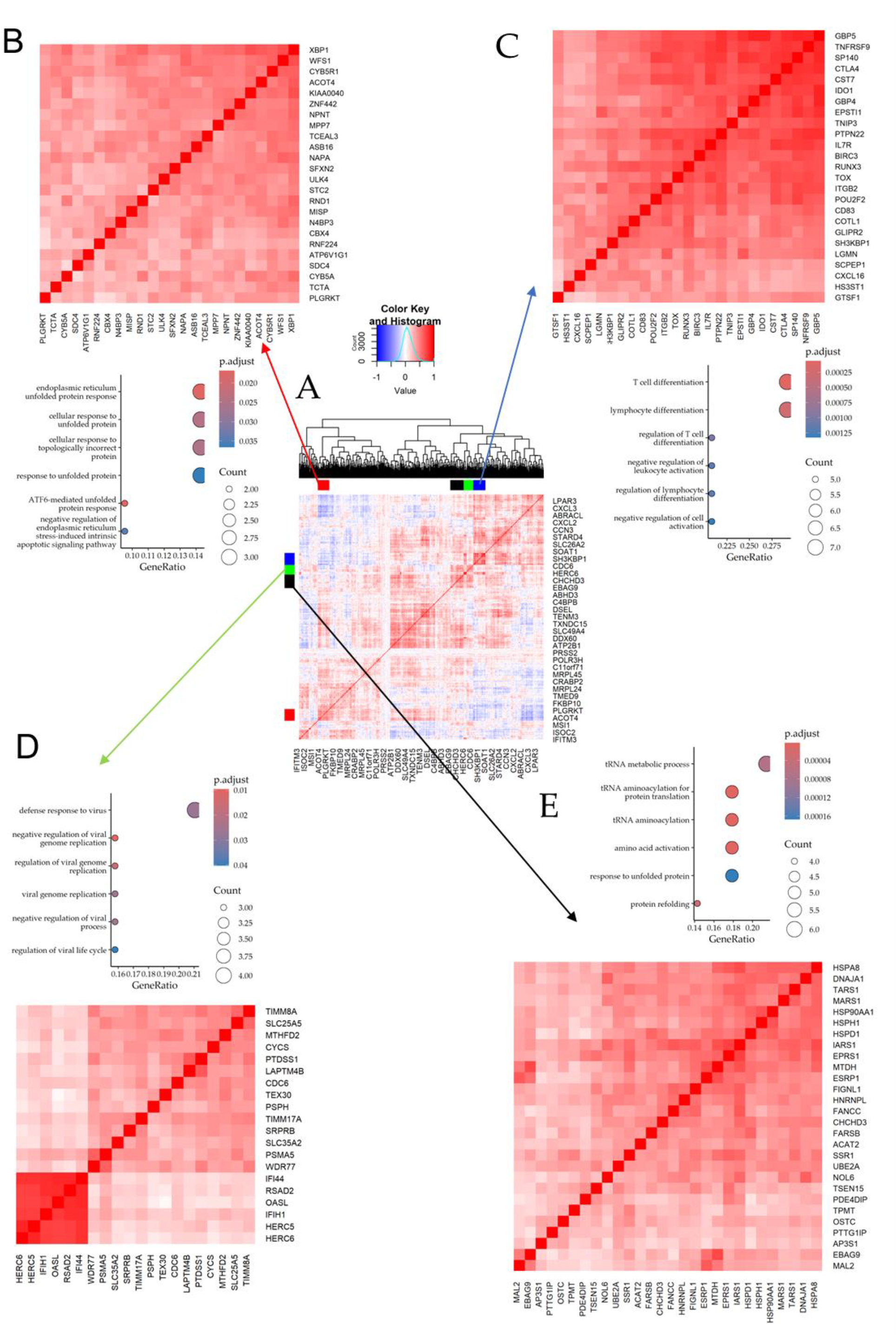
Correlated clusters of oscillatory genes in BRCA show phenotypic relations to misfolded proteins and inflammation. Correlation coefficients of individual Osc^up^ genes show a clustering structure (**A**), with individual subclusters from the hierarchical clustering enriched for (**B**) unfolded protein response, (**C**) T cell differentiation, (**D**) viral response, and (**E**) protein folding.

### Testing gene-wise survival in breast cancer patients

Our data showed that oscillation in the hypoxic milieu is likely to contribute enrichment of many cancer associated phenotypes. Therefore, we asked if hypoxic oscillation induced gene expression may also inform patient survival in breast cancers. The effect of gene expression on the cancer prognosis was calculated using the Cox proportional hazards model. Firstly, the gene expression of each gene was normalized to a z-score to remove artifacts related to the absolute expression ranges of each gene. Statistical tests were conducted for each gene separately. Two versions of the test were employed; survival was predicted using the z-score of the gene expression for each patient (**Figure 5A**), or a binary version wherein the patients were separated into two cohorts of positive and negative z scores (**Figure 5B**). Since the continuous version gave more significant pvalues, the hazard ratio obtained from the z-scores was used for subsequent analysis. **Figure 5C** and **Figure 5D** show an example of an Osc^up^ gene (MCTS1), and an Osc^down^ gene (TP53I11) on survival in breast cancer (BRCA) cohort.

**Figure 5.**
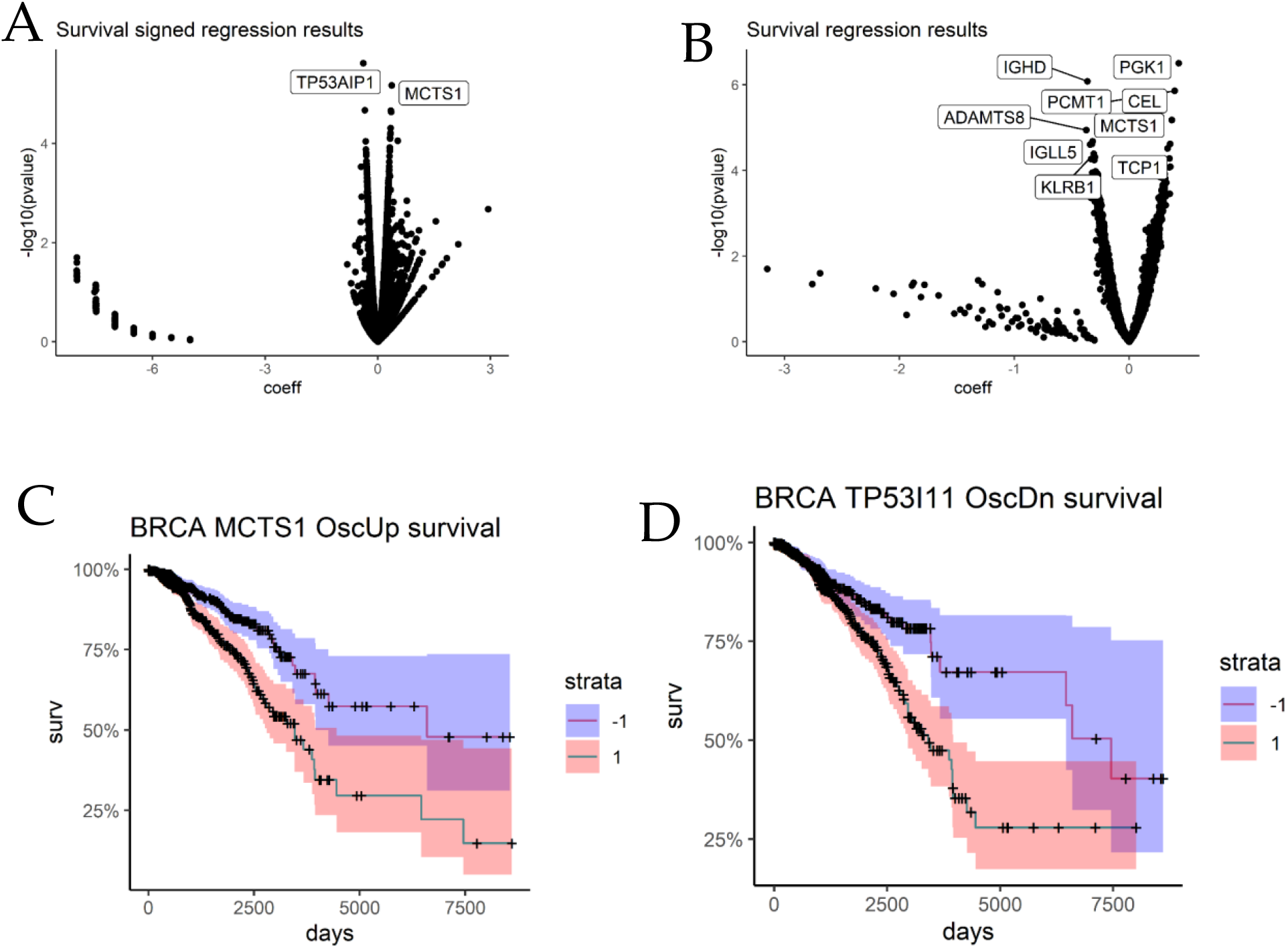
Cox Hazard Ratio calculation for each gene in BRCA. Distribution of the hazard ratios and p-values relating the (**A**) signed and (**B**) raw gene expression z-scores with breast cancer patient survival. Example survival plots for the high and low gene expression cohorts for (**C**) MCTS1, Osc^up^ gene, and (**D**) TP53I11, a Osc^down^ gene.

### Genes upregulated by oscillatory hypoxia prognosticate low survival in breast cancer patients

We ranked genes according to their Cox hazard ratio, and tested the enrichment of Oscup genes. GSEA analysis showed a significant enrichment suggesting that Oscup genes prognosticate low survival in BRCA cancer (**Figure 6A**). We also tested GO enrichment for the leading edge Oscup genes with low survival (low Cox hazard ratio) to explore how oscillatory hypoxia specifically affects gene expression associated with high fatality in BRCA patients. GO analysis again primarily highlighted gene-sets associated with UPR (Figure 6B). The top activated genes in these GOs highlighted many genes associated with malignancy and growth in breast cancers. These included CCN3 encoding a matricellular ligand associated with malignancy in TNBC^17^, UNCB encoding a member of the dependence receptor family associated with high malignancy in breast as well as other cancers, WFS1 encoding wolframin which regulates Ca2+ in ER stress identified as a therapeutic target for colon cancer^18^, NRP1 encoding Neuropilin-1 which regulates breast cancer growth by receptor tyrosine kinase signaling^19^. In addition, were genes encoding members of the heat shock protein family including HSPD1, DNAJA1. We also found genes upregulated in oscillatory hypoxia associated with resistance to chemotherapy which is likely if UPR was upregulated. These included SCG2^20^, and FIGNL1^21^, and NTSR1^22^.

**Figure 6.**
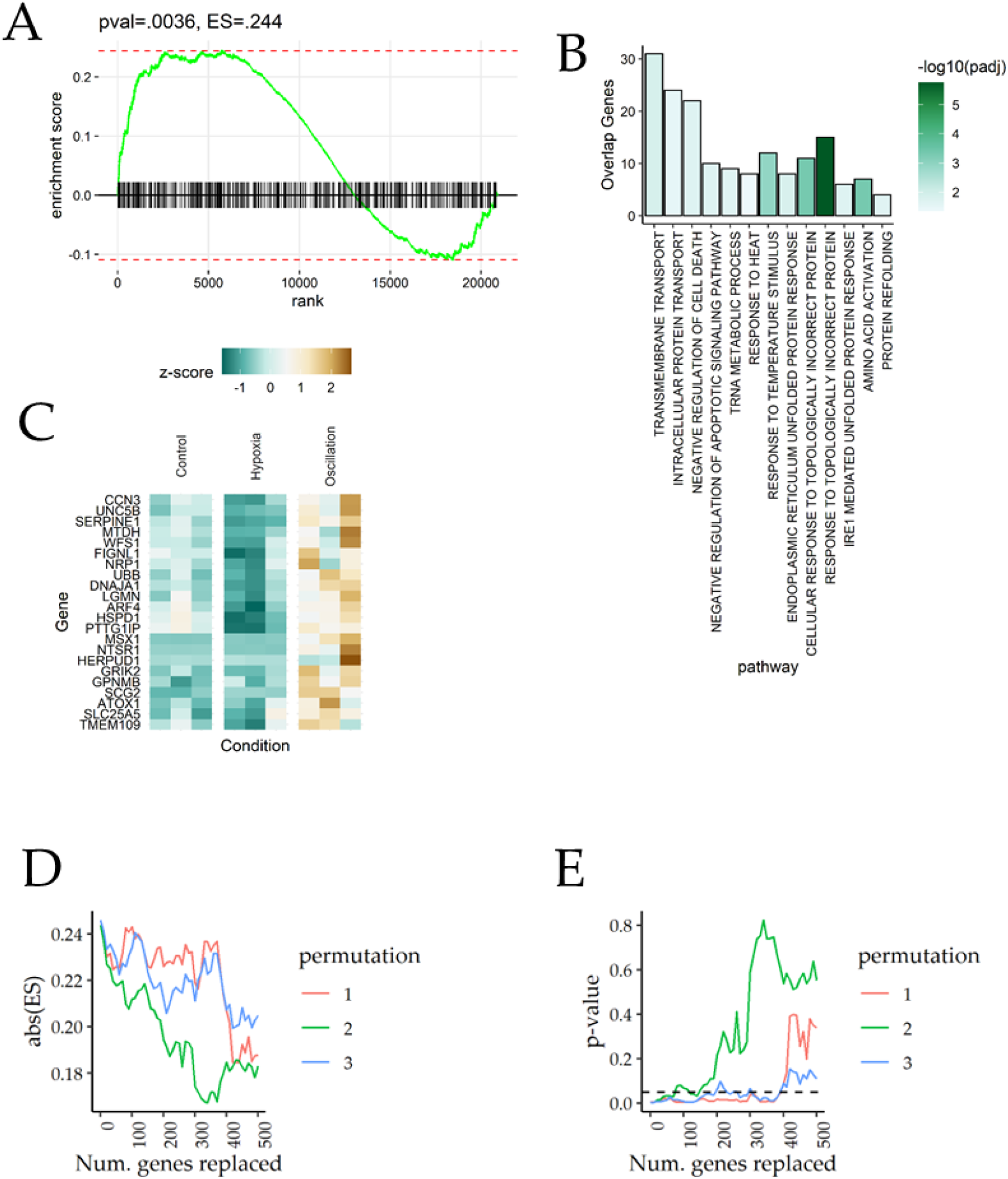
Hypoxic Oscillation specific genes prognosticate low survival in BRCA patients. (**A**) GSEA plot showing the Osc^up^ genes enriched among pro-fatality (i.e., higher hazard ratio) genes, when ranked using hazard ratios. (**B**) Top GO pathways with genes in the leading edge of (**A**) overrepresented in the pathways. (**C**) Heatmap showing the expression of genes in the Negative Regulation of Cell Death pathway that overlap with the leading edge of (**A**). (**D-E**) show the loss of statistical significance and enrichment score with survival ranking when successively replacing the Oscup genes with non-oscillatory genes.

## Discussion

In this work we highlight that dynamics in tumor hypoxia can have a surprising effect on cancer cell gene expression differently from stable hypoxia. It is increasingly recognized that the hypoxic niches in tumors are not uniformly hypoxic in time, and fluctuations in oxygen availability is probably a more common phenomenon than appreciated before. There is yet no consensus about the mechanisms driving fluctuations in hypoxia, although many hypotheses exist, including leaky vasculature, temperature fluctuations etc. Considering how consequential tumor hypoxia is in determining cancer phenotypes, changes in this key microenvironmental factor may itself be consequential. However, little is understood whether hypoxic fluctuations promote or inhibit cancer malignancy, and even less of the mechanisms driving the changed phenotypes.

In addition to oscillation in hypoxia itself, we have previously shown that the transcriptional activity of HIF-1 itself may oscillate in stably high hypoxia. This oscillation is driven by a combination of increased hypoxia driven stability in HIF-1, and increased degradation of HIF-1 by chaperone mediated autophagy. Importantly, we had found that lactate, which is an abundant metabolite in hypoxic tumors drives these oscillations in a small subset of cells. We had shown that emergent oscillations in HIF-1 activity can allow cells to escape cell cycle arrest and continue to proliferate in low oxygen conditions.

In this work, we used MDA-MB-231 cells representing TNBC which is characterized by high intratumoral hypoxia. Gene expression analysis by RNA sequencing in these cells conditioned in normoxia, hypoxia, and an oscillatory hypoxia regimen showed that oscillatory hypoxia had a very specific effect on gene expression, quite different from stable hypoxia. The default expectation is that gene expression in oscillatory hypoxia remains intermediate between normoxia and hypoxia. Although true for most genes, there were also a large number of genes which showed a pattern distinct from hypoxia. Importantly these oscillatory specific genes were highly enriched in most human cancers in The Cancer Genome Atlas (TCGA) database, suggesting that oscillatory hypoxia can specifically upregulate tumor-associated genes, and therefore is likely to be pro-cancer progression factor.

Oscillatory hypoxia occurs in the tumor microenvironment is an emergent phenomenon that arises from the interaction of tumor cells and their metabolic activity, the local microvasculature, and systemic cardiovascular effects. While the results in this paper focus on the effect of oscillatory hypoxia within a cell’s regulatory network, intercellular interactions within a hypoxic microenvironment, stable or fluctuating, may shed further light on the tissue-level manifestation of HIF-1 transcriptional dynamics. Multi-scale modeling of these responses integrating the molecular, cellular, and tissue scales could reveal distinct emergent cellular phenotypes. In future, we envision that functional heterogeneity among tumor cells can be modeled from single cell RNAseq data, and the tumor microenvironment stimulated with different hypoxic dynamics in a multi-scale model consisting of individual cellular automata, interacting with each other.

Categorizing genes increasing in expression in oscillatory hypoxia in (i) genes which showed change in expression even more than in hypoxia, and (ii) those which showed fold change opposite to hypoxia vs normoxia, we found that both the sets were enriched in BRCA cancer samples. Calculating the Cox hazard ratio describing BRCA patient survival for each gene expression, we found that oscillatory genes strongly prognosticate low survival of patients.

A detailed gene-set analysis of these oscillatory genes which also predict low patient survival showed activation of ontologies associated with ion homeostasis, and unfolded protein folding response (UPR). UPR has been implicated in drug resistance in breast cancer and therefore our work hints that fluctuations in hypoxia may cause ER stress, which could promote drug resistance through UPR activation. Overall, our work show that dynamics in hypoxia, and HIF-1 activity may profoundly influence cancer phenotypes differently from stably high HIF-1 activity, both explaining and providing avenues to address yet another mechanism to contribute to heterogeneity in cancer phenotype, and contributing to low patient survival.

## Methods

MDA-MB-231 cells were purchased from ATCC, grown in DMEM/F12 supplemented with 10% FBS (Gibico) and 1% penicillin/streptomycin in a humidified incubator at 37 °C and 5% CO2. Control normoxic cells were maintained in normal atmosphere. The hypoxia cells were incubated at 1% O2 for 48 h in a hypoxia chambers (Embrient Inc) containing 99% N_2_ and 1 % O_2_. Under intermittent hypoxia (Oscillation), cells were subjected to repeated hypoxia (1h, 1% O2)/reoxygenation (30 min, 20% O_2_) cycles for 48h in Coy In vitro Hypoxic Cabinet System (Coy Laboratory Products, Michigan).

Gene expression fold changes between cancer and normal tissue for different cancer datasets were obtained from the Gene Expression Profiling Interactive Analysis (GEPIA)^23^. Patient-wise tumor gene expression and clinical data were extracted from TCGA’s BRCA project (https://portal.gdc.cancer.gov/projects/TCGA-BRCA)^11^.

Student’s t-tests were conducted for differences in the mean fold changes of ELI genes vs. all other genes. Statistical significance (p-values) and effect sizes (differences in means) are reported in the figures.

The effect of gene expression on the cancer prognosis was calculated using the Cox proportional hazards model. Firstly, the gene expression of each gene was normalized to a z-score to remove artifacts related to the absolute expression ranges of each gene. Statistical tests were conducted for each gene separately. Two versions of the test were employed; survival was predicted using the z-score of the gene expression for each patient, or a binary version wherein the patients were separated into two cohorts of positive and negative z scores. Since the continuous version gave more significant p-values, the hazard ratio obtained from the z-scores was used for subsequent analysis. To find the relationship between the Osc^up^ and Osc^down^ genes, genes were ranked according to their hazard ratios.

All Gene Set Enrichment Analysis (GSEA) followed the standard Kolmogorov–Smirnov tests, and were conducted using the fgsea package^24^, and visualized using the method of Subramaniam et al^25^.

## Acknowledgements

Funding was provided by National Cancer Institute R37 grant (R37 CA248161/CA/NCI NIH HHS/United States). The results shown here are in part based upon data generated by the TCGA Research Network: https://www.cancer.gov/tcga^11^.

## References

1. Weidemann, A. & Johnson, R. S. Biology of HIF-1α. Cell Death and Differentiation vol. 15 621–627 Preprint at 10.1038/cdd.2008.12 (2008).

2. Semenza, G. L. Hypoxia-Inducible Factors in Physiology and Medicine. Cell 148, 399–408 (2012).

3. Semenza, G. L. Hypoxia-inducible factor 1 (HIF-1) pathway. Sci STKE 2007, (2007).

4. Wohlrab, P. et al. Oxygen conditions oscillating between hypoxia and hyperoxia induce different effects in the pulmonary endothelium compared to constant oxygen conditions. Physiol Rep 9, (2021).

5. Mironov, S. L. & Richter, D. W. Oscillations and hypoxic changes of mitochondrial variables in neurons of the brainstem respiratory centre of mice. J Physiol 533, 227 (2001).

6. Milotti, E., Stella, S. & Chignola, R. Pulsation-limited oxygen diffusion in the tumour microenvironment. Scientific Reports 2017 7:1 7, 1–11 (2017).

7. Kshitiz et al. Lactate-dependent chaperone-mediated autophagy induces oscillatory HIF-1α activity promoting proliferation of hypoxic cells. Cell Syst 13, 1048–1064.e7 (2022).

8. Bagnall, J. et al. Tight Control of Hypoxia-inducible Factor-α Transient Dynamics Is Essential for Cell Survival in Hypoxia. Journal of Biological Chemistry 289, 5549–5564 (2014).

9. Colegio, O. R. et al. Functional polarization of tumour-associated macrophages by tumour-derived lactic acid. Nature 513, 559 (2014).

10. Movafagh, S., Crook, S. & Vo, K. Regulation of hypoxia-inducible factor-1α by reactive oxygen species: new developments in an old debate. J Cell Biochem 116, 696–703 (2015).

11. Liu, J. et al. An Integrated TCGA Pan-Cancer Clinical Data Resource to Drive High-Quality Survival Outcome Analytics. Cell 173, 400–416.e11 (2018).

12. Schwab, A. & Stock, C. Ion channels and transporters in tumour cell migration and invasion. Philos Trans R Soc Lond B Biol Sci 369, (2014).

13. Jeong, J. & Eide, D. J. The SLC39 family of zinc transporters. Mol Aspects Med 34, 612 (2013).

14. Forte, M. et al. An interplay between UCP2 and ROS protects cells from high-salt-induced injury through autophagy stimulation. Cell Death & Disease 2021 12:10 12, 1–12 (2021).

15. Son, S. et al. CCN3/NOV promotes metastasis and tumor progression via GPNMB-induced EGFR activation in triple-negative breast cancer. Cell Death & Disease 2023 14:2 14, 1–16 (2023).

16. Yang, X. et al. System analysis based on the ER stress-related genes identifies WFS1 as a novel therapy target for colon cancer. Aging 14, 9243–9263 (2022).

17. Abdullah, A. et al. Epigenetic targeting of neuropilin-1 prevents bypass signaling in drug-resistant breast cancer. Oncogene 2020 40:2 40, 322–333 (2020).

18. Weng, S. et al. SCG2: A Prognostic Marker That Pinpoints Chemotherapy and Immunotherapy in Colorectal Cancer. Front Immunol 13, 873871 (2022).

19. Meng, C. et al. FIGNL1 is a potential biomarker of cisplatin resistance in non-small cell lung cancer. International Journal of Biological Markers (2022) doi:10.1177/03936155221110249/ASSET/IMAGES/LARGE/10.1177_03936155221110249-FIG6.JPEG.

20. Wu, Z., Martinez-Fong, D., Trédaniel, J. & Forgez, P. Neurotensin and its high affinity receptor 1 as a potential pharmacological target in cancer therapy. Front Endocrinol (Lausanne) 3, 39278 (2013).

21. Tang, Z., Kang, B., Li, C., Chen, T. & Zhang, Z. GEPIA2: an enhanced web server for large-scale expression profiling and interactive analysis. Nucleic Acids Res 47, W556–W560 (2019).

22. Korotkevich, G. et al. Fast gene set enrichment analysis. bioRxiv 060012 (2021) doi:10.1101/060012.

23. Subramanian, A. et al. Gene set enrichment analysis: A knowledge-based approach for interpreting genome-wide expression profiles. Proc Natl Acad Sci U S A 102, 15545–15550 (2005).

## References

1 Weidemann, A. & Johnson, R. S. Biology of HIF-1alpha. Cell Death Differ 15, 621–627 (2008). 10.1038/cdd.2008.12

2 Semenza, G. L. Hypoxia-inducible factor 1 (HIF-1) pathway. Sci STKE 2007, cm8 (2007). 10.1126/stke.4072007cm8

3 Dengler, V. L., Galbraith, M. & Espinosa, J. M. Transcriptional regulation by hypoxia inducible factors. Crit Rev Biochem Mol Biol 49, 1–15 (2014). 10.3109/10409238.2013.838205

4 Wohlrab, P. et al. Oxygen conditions oscillating between hypoxia and hyperoxia induce different effects in the pulmonary endothelium compared to constant oxygen conditions. Physiol Rep 9, e14590 (2021). 10.14814/phy2.14590

5 Mironov, S. L. & Richter, D. W. Oscillations and hypoxic changes of mitochondrial variables in neurons of the brainstem respiratory centre of mice. J Physiol 533, 227–236 (2001). 10.1111/j.1469-7793.2001.0227b.x

6 Milotti, E., Stella, S. & Chignola, R. Pulsation-limited oxygen diffusion in the tumour microenvironment. Sci Rep 7, 39762 (2017). 10.1038/srep39762

7 Kshitiz et al. Lactate-dependent chaperone-mediated autophagy induces oscillatory HIF-1alpha activity promoting proliferation of hypoxic cells. Cell Syst 13, 1048–1064 e1047 (2022). 10.1016/j.cels.2022.11.003

8 Bagnall, J. et al. Tight control of hypoxia-inducible factor-alpha transient dynamics is essential for cell survival in hypoxia. J Biol Chem 289, 5549–5564 (2014). 10.1074/jbc.M113.500405

9 Colegio, O. R. et al. Functional polarization of tumour-associated macrophages by tumour-derived lactic acid. Nature 513, 559–563 (2014). 10.1038/nature13490

10 Movafagh, S., Crook, S. & Vo, K. Regulation of hypoxia-inducible factor-1a by reactive oxygen species: new developments in an old debate. J Cell Biochem 116, 696–703 (2015). 10.1002/jcb.25074

11 Liu, J. et al. An Integrated TCGA Pan-Cancer Clinical Data Resource to Drive High-Quality Survival Outcome Analytics. Cell 173, 400–416 e411 (2018). 10.1016/j.cell.2018.02.052

12 Schwab, A. & Stock, C. Ion channels and transporters in tumour cell migration and invasion. Philos Trans R Soc Lond B Biol Sci 369, 20130102 (2014). 10.1098/rstb.2013.0102

13 Jeong, J. & Eide, D. J. The SLC39 family of zinc transporters. Mol Aspects Med 34, 612–619 (2013). 10.1016/j.mam.2012.05.011

14 SenGupta, S. et al. Triple-Negative Breast Cancer Cells Recruit Neutrophils by Secreting TGF-beta and CXCR2 Ligands. Front Immunol 12, 659996 (2021). 10.3389/fimmu.2021.659996

15 Ruffell, B. et al. Leukocyte composition of human breast cancer. Proc Natl Acad Sci U S A 109, 2796–2801 (2012). 10.1073/pnas.1104303108

16 Forte, M. et al. An interplay between UCP2 and ROS protects cells from high-salt-induced injury through autophagy stimulation. Cell Death Dis 12, 919 (2021). 10.1038/s41419-021-04188-4

17 Son, S. et al. CCN3/NOV promotes metastasis and tumor progression via GPNMB-induced EGFR activation in triple-negative breast cancer. Cell Death Dis 14, 81 (2023). 10.1038/s41419-023-05608-3

18 Yang, X. et al. System analysis based on the ER stress-related genes identifies WFS1 as a novel therapy target for colon cancer. Aging (Albany NY) 14, 9243–9263 (2022). 10.18632/aging.204404

19 Abdullah, A. et al. Epigenetic targeting of neuropilin-1 prevents bypass signaling in drug-resistant breast cancer. Oncogene 40, 322–333 (2021). 10.1038/s41388-020-01530-6

20 Weng, S. et al. SCG2: A Prognostic Marker That Pinpoints Chemotherapy and Immunotherapy in Colorectal Cancer. Front Immunol 13, 873871 (2022). 10.3389/fimmu.2022.873871

21 Meng, C. et al. FIGNL1 is a potential biomarker of cisplatin resistance in non-small cell lung cancer. Int J Biol Markers 37, 260–269 (2022). 10.1177/03936155221110249

22 Wu, Z., Martinez-Fong, D., Tredaniel, J. & Forgez, P. Neurotensin and its high affinity receptor 1 as a potential pharmacological target in cancer therapy. Front Endocrinol (Lausanne) 3, 184 (2012). 10.3389/fendo.2012.00184

23 Tang, Z., Kang, B., Li, C., Chen, T. & Zhang, Z. GEPIA2: an enhanced web server for large-scale expression profiling and interactive analysis. Nucleic Acids Res 47, W556–W560 (2019). 10.1093/nar/gkz430

24 Gennady Korotkevich, V. S., Nikolay Budin, Boris Shpak, Maxim N. Artyomov, Alexey Sergushichev. Fast gene set enrichment analysis. bioRxiv 060012 (2021).

25 Subramanian, A. et al. Gene set enrichment analysis: a knowledge-based approach for interpreting genome-wide expression profiles. Proc Natl Acad Sci U S A 102, 15545–15550 (2005). 10.1073/pnas.0506580102

